# Quantitative solid-phase assay to measure deoxynucleoside triphosphate pools

**DOI:** 10.1101/270454

**Authors:** Juan Cruz Landoni, Liya Wang, Anu Suomalainen

**Author notes:** To whom correspondence should be addressed. Prof. Anu Suomalainen, Biomedicum-Helsinki, Molecular Neurology Research Program, Haartmaninkatu 8, University of Helsinki, 00290 Helsinki, Finland.

## Abstract

Deoxyribonucleoside triphosphates (dNTPs) are the reduced nucleotides used as the building blocks and energy source for DNA replication and maintenance in all living systems. They are present in highly regulated amounts and ratios in the cell, and their balance has been implicated in the most important cell processes, from determining the fidelity of DNA replication to affecting cell fate. Furthermore, many cancer drugs target biosynthetic enzymes in dNTP metabolism, and mutations in genes directly or indirectly affecting these pathways are the cause of devastating diseases. The accurate and systematic measurement of these pools is key to understand the mechanisms behind these diseases and their treatment. We present a new method for measuring dNTP pools from biological samples, utilising the current state-of-the-art polymerase method, modified to a solid-phase setting and optimised for larger scale measurements.

## INTRODUCTION

Deoxyribonucleic acid (DNA) carries the genetic information of all known living organisms, and its synthesis utilises deoxynucleoside triphosphates (dNTPs), both as building blocks and as energy source for the polymerisation reaction. dNTP pools depend on cell cycle and cell and tissue type, with proliferating cells typically having higher pools than post-mitotic cells (1). dNTPs are typically present at very low concentrations, which are tightly regulated via the expression, regulation and localisation of their biosynthetic and catabolic enzymes (2–4). These enzymes form a complex and highly regulated network involving, in animal cells, the nuclear, mitochondrial and cytoplasmic compartments. These interlinked pathways include the biosynthetic *de novo* pathway and the parallel cytoplasmic and mitochondrial salvage pathways (5, 6).

The absolute amounts and ratios of the different dNTPs in time and space have been shown to be critical for a wide range of cellular processes, including DNA replication fidelity and mutagenesis (7, 8), DNA repair, cell cycle progression and regulation, mitochondrial DNA (mtDNA) maintenance function and oncogenic and apoptotic processes (9). Because of the mutagenic effects of imbalanced dNTP pools, they have been studied in detail in the context of cancer biology. However, accumulating data points to tight co-regulation of the mitochondrial and cytoplasmic pools, evidenced by defects in the core synthetic enzymes causing depletion or mutagenesis of mtDNA (10, 11). These disorders present varying tissue specificity and different ages of onset, typically with multi-systemic devastating consequences (12–14). Furthermore, disorders of mitochondrial DNA maintenance can secondarily affect the cytoplasmic dNTP pool balance (15, 16), underscoring the intimate crosstalk of the different compartmentalised dNTP pools. Preclinical evidence suggest that some mtDNA instability disorders could be treated with nucleosides (17–19). However, knowledge of the regulation of the dNTP pools in pathological metabolic states or upon supplementation therapies is still scarce, and sensitive methodology is needed.

The most widely used approach to measure dNTP concentrations from biological samples is based on a polymerase assay using radioactively labelled substrate, first proposed in the late 1960s (20) and further applied and developed two decades later (21). This traditional method is laborious and multistep, challenging its sensitivity. After different modifications and the replacement of the Klenow fragment for a thermostable polymerase to improve ribonucleotide discrimination (22), the most recent update for this method was developed for the measurement of mitochondrial dNTP pools (23). The assay relies on the specificity of a polymerase to utilise the dNTPs present in the sample, and bind them on a designed template, together with a different radio-labelled dNTP which will be incorporated proportionally. The amount of the radioactive labelled products (newly synthesised oligonucleotides), together with a standard series of known dNTP concentrations, provides a quantitative result. However, the field has been hampered by the recent restricted commercial availability of specific materials needed for this protocol, motivating method development.

We present here the adaptation of the dNTP measurement into a solid-phase format, utilising combined knowledge from the conventional polymerase-based dNTP measurement and the “solid-phase mini-sequencing” principle of single-nucleotide detection (24, 25). This allows a microtiter-plate-based, automatable approach, with an accurate measurement of a large number of samples, improving efficiency, safety and accuracy of the methodology.

## MATERIALS AND METHODS

**Fig. 1A** shows an example of the reaction in the setting of the present method, being used to measure deoxycytidine triphosphate (dCTP).

**Figure 1:**
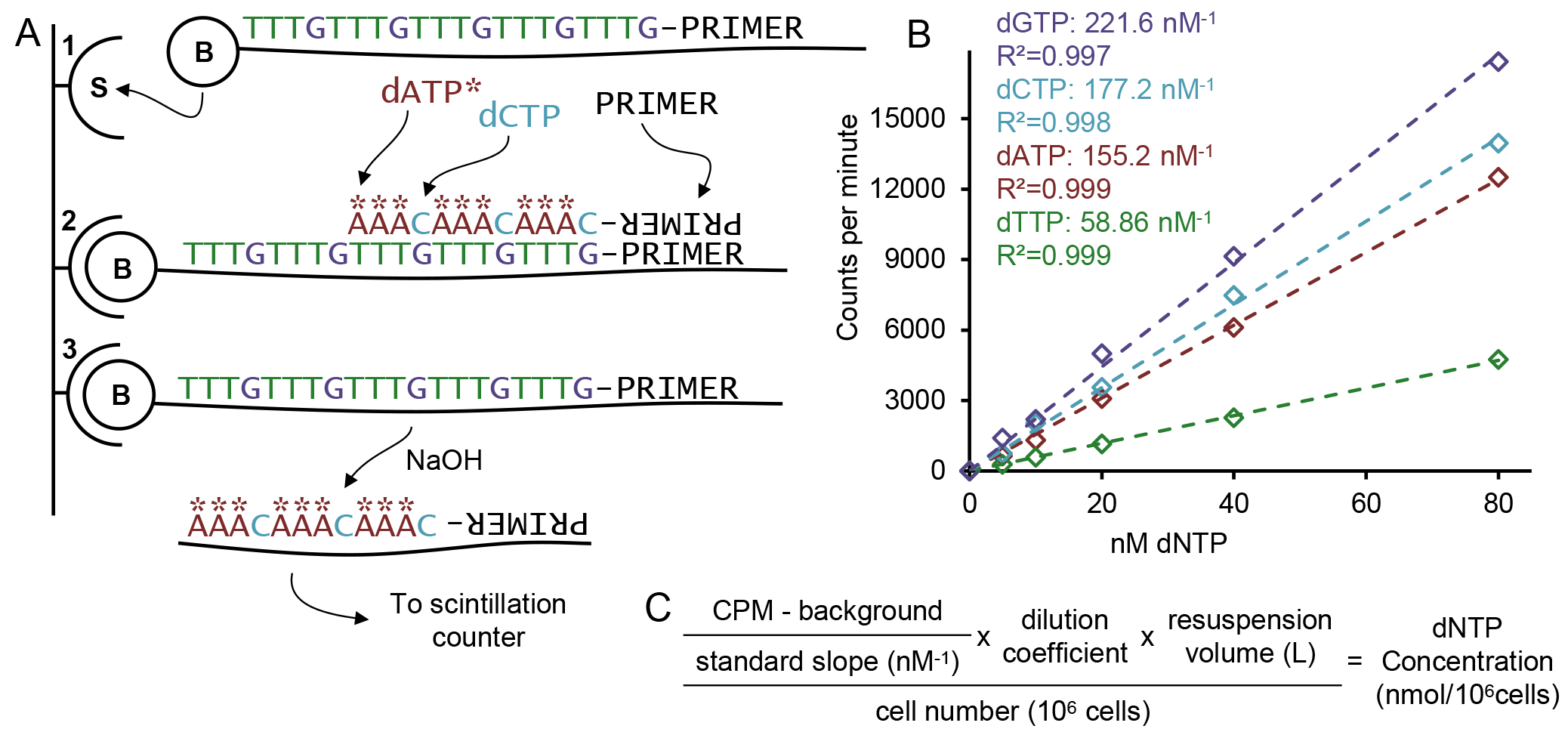
(A) Schematic description of the reaction principle using dCTP measurement as an example. (A1) Affinity capture of biotinylated oligonucleotide OligoC to streptavidin-coated plate. (A2) Polymerisation: priming of the template, followed by the proportional incorporation of the measured nucleotide (dCTP) and the radiolabelled nucleotide (dATP*). (A3) Alkaline release of the labelled chain and radioactive count. (B) Example of standard curves and slopes to be used in the data analysis, with the regression’s calculated R^2^ for each nucleotide measurement. (C) Equation for the analysis of the counts per minute (CPM) obtained from the procedure, utilising the resulting slope from the standards and the blank, as well as the recorded dilution factors and cell number to obtain the concentration of the nucleotide per million cells.

### Biotinylated oligonucleotides and primer

**Table 1** below shows the sequences of the primer and the oligonucleotides used in the reaction. The templates are labelled “Oligo”+N, with N being the specific dNTP to be measured with that template and the primer being common to all reactions.

**Table 1.**
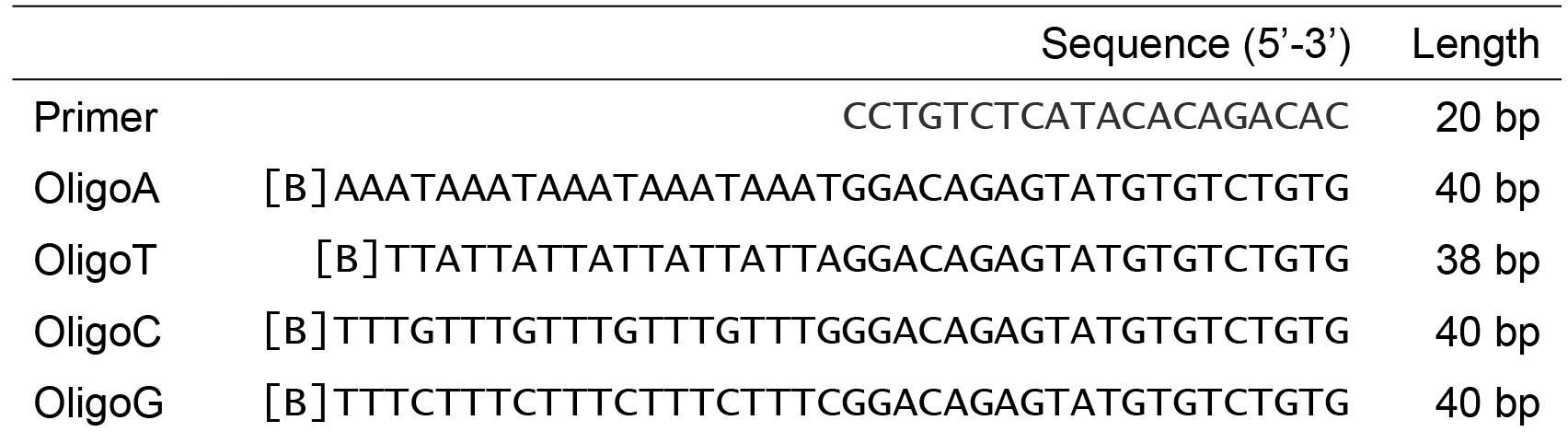

The biotin-labelled oligonucleotides should be HPLC-purified and, as well as the primer, diluted to 5 μM concentration for the reaction, aliquoted and stored at −20°C.

### Reagents and equipment

(In parenthesis, the name of the product/company used in the set-up)

– 60% methanol in water, kept at −20°C.
– 50 mM NaOH (made fresh every 4 weeks).
– Thermostable DNA polymerase (DyNAzyme II DNA Polymerase by ThermoFisher Scientific), with its 10x optimised Enzyme Buffer, stored at −20°C.
– 0.5 M dithiothreitol (DTT) in small aliquots stored at −20°C (single use because of instability).
– dNTP mix stock, 40 mM dNTPs (10 mM each) (Bioline), and series dilutions until 1 μM stored at −20°C.
– [8-^3^H(N)]-Deoxyadenosine 5’-triphosphate Tetrasodium Salt in 1:1 ethanol:water mixture (PerkinElmer), stored at −20°C.
– [Methyl-^3^H]-Deoxythymidine 5`-Triphosphate Tetrasodium Salt in 1:1 ethanol:water mixture (PerkinElmer), stored at −20°C.
– Phosphate buffered Saline (PBS).
– TWEEN^®^ 20 (Amresco).
– TENT solution (40 mM Tris-HCl, 1mM EDTA, 50mM NaCl, 0.1% TWEEN^®^ 20, pH 8.0-8.8).
– Biotinylated oligonucleotides and primer (Sigma Aldrich), stock and dilutions in −20°C.
– Streptavidin coated 96-well plate (BioBind Streptavidin Strip Assembled Solid by Thermo Scientific).
– Ultima Gold™ liquid scintillation cocktail for aqueous and non-aqueous samples (PerkinElmer).
– Scintillation vials and beta counter (MicroBeta^2^ by PerkinElmer).
– Microplate washer (Wellwash™ by Thermo Labsystems).
– Speed vacuum concentrator.
– Countess automated cell counter (Invitrogen).

### dNTP isolation

1. Culture cells in Petri dishes to an ideal minimum of 1 ⨯10^6^ cells. As dNTPs vary greatly between dividing and non-dividing cells and throughout the cell cycle, plan to harvest the cells in similar confluency and cell cycle stage. Wash the cells carefully with cold PBS and detach by trypsinisation.
2. For normalisation, count the cell concentration in the suspension. Then pellet the cells in a microcentrifuge at 250·*g*, and store the pellet at −80 °C.
3. For the isolation, add 1.5 ml of cold (−20 °C) 60 % methanol in water and vortex thoroughly. Then incubate for at least 1 hour at −80 ° C.
4. Pellet the insoluble cell contents for 15 minutes at maximum speed (20 000·*g* at 4 ° C); incubate the samples at 95 °C for 3 minutes, cool down on ice, and pellet again similarly.
5. Collect the supernatant and transfer it to a new tube. Desiccate the supernatant with a speed vacuum concentrator until fully dry.
6. Store the solid extract at −80 °C until analysis.

### Solid-phase radioactive polymerase reaction

1. *Affinity capture of oligonucleotides:* Each dNTP will be measured in a separate well. Mark the plates carefully for the four dNTP reaction, for the standard series and sample replicates. Each measurement replicate requires four wells (one per dNTP), and it is recommended to have at least duplicates at different dilutions to confirm linearity. The standard series of six concentrations will use 24 wells. Transfer 2.5 μl of the specific template oligonucleotide into each streptavidin-coated well, and add 47.5 μl of a 0.1% TWEEN^®^ in PBS solution. Incubate the plate at 37 °C for 1.5 h with gentle shaking.
2. Discard the liquid from the wells, and wash the wells thoroughly four times at room temperature with 200 μl of TENT solution. An automated plate washer can be used. Tap the wells dry against tissue papers after the washes.
3. *Standards:* Prepare a standard dilution series of a commercial dNTP mixture, with the following concentrations: 80 nM, 40 nM, 20 nM, 10 nM, 5 nM, and a water blank.
4. *Sample preparation:* Dissolve the frozen solid nucleotide extract in cold sterile water (normally 100 μl, modify according to the cell number), vortex thoroughly and keep on ice for 10 minutes.
5. Prepare replicates of the different dilution coefficients (normally 1:5 and 1:10) for each sample to be measured, in a volume sufficient for four reactions per dilution (minimum 50 μl).
6. *Reaction:* Prepare master-mixes for each nucleotide to be measured according to the **Table 3**. [^3^H]-dATP is used for measuring dTTP, dCTP and dGTP (the wells with OligoT, OligoC and OligoG) in the sample, while [^3^H]-dTTP is used for the dATP (OligoA) reaction mixture. Since the radio-labelled dNTPs are stored in ethanol:water mixture, the molar concentration of each batch should be calculated from the activity per volume and specific radioactivity, and the solvent of the required amount evaporated with a speed-vacuum concentrator, as recommended by (23).
7. Incubate the plate at 55°C for 1 hour, with a flotation device in a water bath.
8. Discard the contents of the wells and perform the washing process as in step 2.
9. Release the newly synthesised labelled strand by adding 60 μl of 50 mM NaOH to each well and incubate for 3 min at room temperature.
10. Transfer the eluted DNA to scintillation vials into 3 ml of scintillation cocktail and measure the radioactivity in a beta counter, with 1 minute counting time per sample.
11. Data analysis: After exporting the data, analyse the counts-per-minute (CPM) values for each nucleotide independently. Subtract the blank value from all samples measuring that nucleotide. Next, generate a linear standard curve from the origin (**Fig 1B**), and use the regression slope to calculate the concentration of the measured replicates based on their CPM-values. Values outside the range of standards should be omitted and re-measured at a different dilution. Then use the cell number and dilution factors utilised to calculate the amount of dNTP per number of cells (**Fig 1C**).

**Table 2.**

| | Volume ($\mu\text{l}$) | Final concentration |
| --- | --- | --- |
| PBS/TWEEN® solution | 47,5 |  |
| 5 $\mu\text{M}$ oligonucleotide | 2,5 | 0,25 $\mu\text{M}$ |
| Total volume | 50 |  |

**Table 3.**

| | Volume (μl) | Final concentration |
| --- | --- | --- |
| 10x polymerase buffer | 5 | 1x |
| 0.5 M DTT | 0.5 | 5 mM |
| 15 μM radiolabelled dNTP | * | 0.75 μM |
| 5 μM primer | 2.5 | 0.25 μM |
| Thermostable polymerase | ** | 0.025 U/μl |
| Sample | 12.5 |  |
| Water | 28.5 |  |
| Total volume | ~50 |  |

\* calculated for each batch and desiccated in the tube.
\*\* calculated from concentration of polymerase.

## RESULTS

The assay was performed on an array of different cell lines. The standard curves show consistently strong linearity within the measurement range, with R^2^ values for the regression around 0.99 (**Fig. 1B**). Each line or cell type presented a particular dNTP pool profile (**Fig. 2A**). The measurements show good reproducibility with same samples measured on different days (*SH-SY5Y* 1-3), as well as with independently cultured and differentiated samples from a same cell line (*Myotubes* 1-2) (**Fig. 2B**).

Finally, in order to compare the performance of the methodology with the traditional one, we measured identical aliquots of mouse bone marrow extracts with a modified version of the protocol in (23) for whole cell lysates, and with the present method. The resulting values (**Fig. 2C-D**) show very similar dNTP ratios, with an increased absolute value, particularly strong concerning the measurement of dTTP.

**Figure 2:**
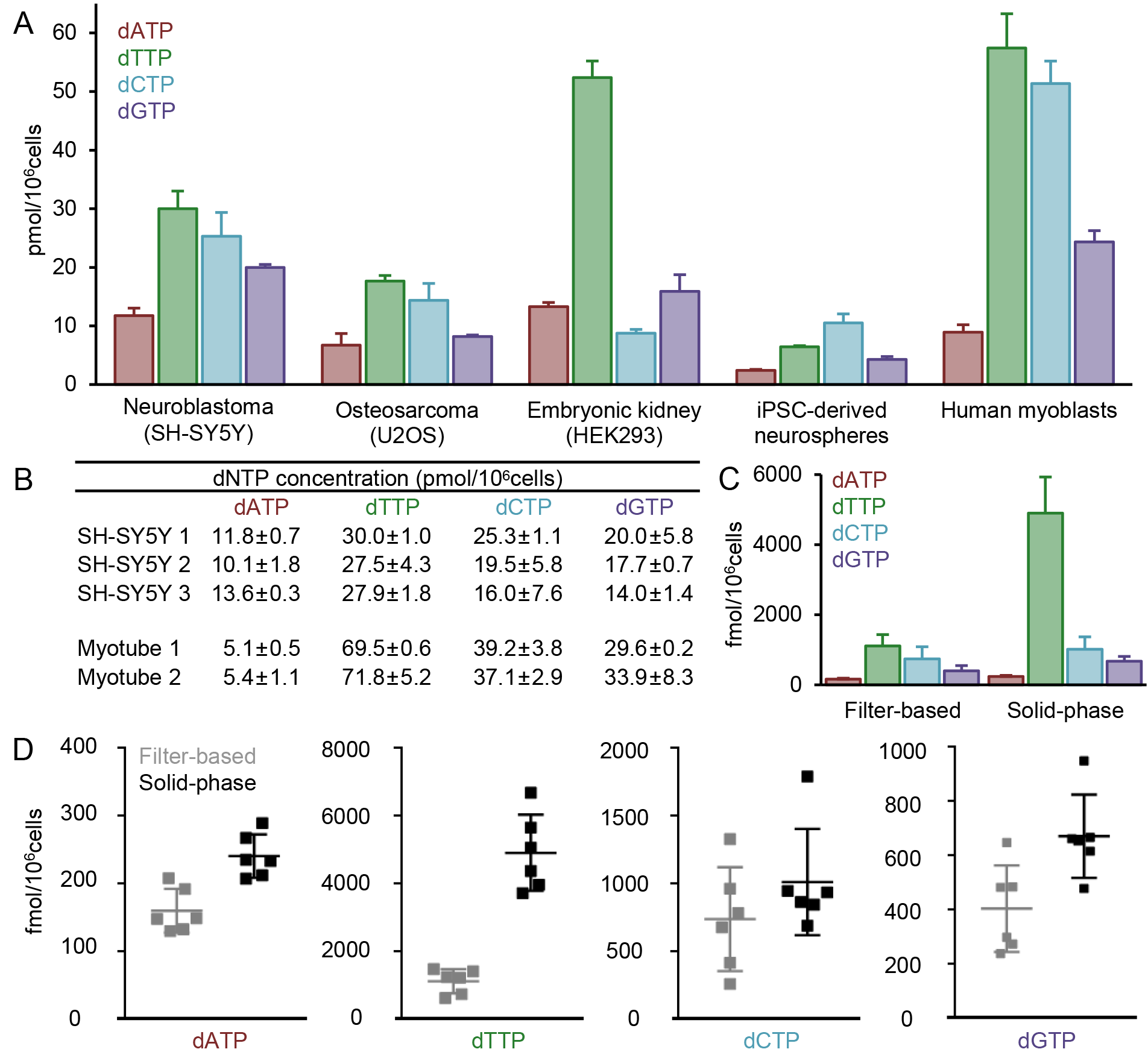
(A) dNTP concentration profiles from an array of cell lines. (B) Concentrations obtained from the measurement of the same sample on different days (*SH-SY5Y* 1 vs. 2) and from independently cultured samples of the same cell line (*SH-SY5Y* 1-2 vs. 3 and *Myotubes* 1 vs 2). (CD) Comparison between the dNTP concentration values of identical isolates of mouse bone marrow cells (n=6) measured with the traditional filter-based protocol or with the present solid-phase methodology.

## DISCUSSION

We report here a development to the current state-of-the-art methodology for quantitative measurement of the four canonical deoxynucleoside triphosphates from biological samples. The method allows the performance of the measurement in a solid-phase multi-well setting, and the microtiter-plate format allows automation of the pipetting and washing steps, as well as circumventing time-consuming primer-preparation and filter-based steps from the previous methods (23). Furthermore, it allows utilisation of optimised reaction buffers and template sequences. All these features dramatically reduce the required analysis steps, time and manual work required, diminishing the potential sources of error. Importantly, it also increases method safety, reducing direct contact with hazardous compounds.

The current shortcoming of the methodology is the increase of the reaction volume due to the nature of the well, leading to a larger requirement of sample and reagents. Furthermore, the normalisation strategy is a challenge in the field (1), due to the inaccuracy of cell counting and difficulties comparing results from different publications. The most widely used normalisation method is cell number, although other strategies are possible, such as utilising the absorbance of an alkaline lysate, proportional to cell mass (26). The consistency of the results obtained by cell number normalisation lead us to utilise it due to its simplicity, but a more sensitive approach should be considered for more challenging biological samples, such as protein or DNA content measurement, or evaluation of cell volume for molar concentration presentation.

Nucleotide metabolism is one of the main targets of chemotherapeutic drugs in cancer, despite the fact that current understanding on nucleotide metabolism and tissue specific features of dNTP regulation is superficial. Furthermore, mitochondrial dysfunction has turned out to be a common cause of inherited degenerative conditions, also causing rare and devastating mitochondrial disorders in children, where nucleoside therapies are emerging as a potential therapeutic option (12). Our sensitive, quantitative and efficient method should be a highly useful tool to provide evidence of dNTP metabolism in different biological samples, in pathological states and in treatment of cancer and metabolic diseases.

